# Vector Trace cells in the Subiculum of the Hippocampal formation

**DOI:** 10.1101/805242

**Authors:** Steven Poulter, Sang Ah Lee, James Dachtler, Thomas J. Wills, Colin Lever

## Abstract

Successfully navigating in physical or semantic space requires a neural representation of allocentric (map-based) vectors to boundaries, objects, and goals. Cognitive processes such as path-planning and imagination entail recall of vector representations, but evidence of neuron-level memory for allocentric vectors has been lacking. Here we describe a novel neuron type (Vector Trace cell, VTC) whose firing generates a new vector field when a cue is encountered, and also a ‘trace’ version of that field for hours after cue removal. VTCs are concentrated in subiculum distal to CA1. Compared to non-trace cells, VTCs fire at further distances from cues, and exhibit earlier-going shifts in preferred theta phase in response to newly introduced cues, demonstrating a theta-linked neural substrate for memory encoding. VTCs suggest a vector-based model of computing spatial relationships between an agent and multiple spatial objects, or between different objects, freed from the constraints of direct perception of those objects.

## Introduction

Neurons in the hippocampal formation represent an organism’s allocentric location and heading ^1–6^, and the fundamental coding underlying these spatial representations, for instance involving the theta oscillation, likely support planning, imagination, and memory beyond the purely spatial domain ^4, 7–9^. Spatial coding may be vector-based, such that a neuron fires at a particular allocentric distance and direction from an environmental boundary or object ^10–13^, and path-planning and imagination often entail the recall of vector representations of environmental cues ^8, 14–16^.

Here, we examined vector-based representations in the subiculum. The subiculum is a major output region of the hippocampal formation ^17–20^. The subiculum is known to contain vector representations ^11^, is implicated in memory retrieval ^21–23^, and has recently been identified as likely the hippocampal component of the default mode network ^24^. This could suggest a wide role for subicular vector-based representations in directing navigation and memory-based cognition, consistent with models of spatial memory and imagery ^8^.

Accordingly, we exposed rats to a range of cues differing in size, shape and sensory properties, and tested for memory-based responses in subicular neurons following cue removal.

## Results

### Cue responsive cells: Defining Vector Trace cells (VTCs) and non-trace cells

Fig. 1A shows a schematic of the experimental manipulations. The dataset comprised only subicular cue-responsive neurons which showed spatial tuning to environmental boundaries ^11^ and inserted cues. Responsiveness to inserted cues was defined by the appearance of a new firing field (‘cue-field’) in the cue trial (198 cells passing this criterion: see Methods). We quantified the strength of memory-based firing in the region of the cue-field, in the post-cue trial using two measures: 1) a ‘Trace score’, which measured the strength of firing in the cue-field region, in the post-cue trial; 2) an ‘Overlap score’, which measured whether firing in the post-cue trial, outside of the wall field (termed the ‘post-cue field’) was spatially overlapping with the cue field region. (Fig. 1B, rightmost column, Fig.1C; see Methods). Trace and Overlap scores together define a group of neurons which show memory-based firing persistence, specifically in the region of the cue-field (firing hereafter referred to as the ‘Trace field’). For further analysis, those neurons with Trace score>0.2 and Overlap>0.4 were defined as showing trace responses, here termed ‘Vector Trace cells’ (VTCs) (70/198; Fig.1C; Fig.S1 shows co-recorded examples of both cell types: VTCs and non-trace cells). The proportion of neurons classified as VTCs was significantly greater than would be expected from spatially random localization of post-cue firing (Z-test for proportions, Z=9.3, p=1.3×10^19^, see Methods).

**Fig. 1.**
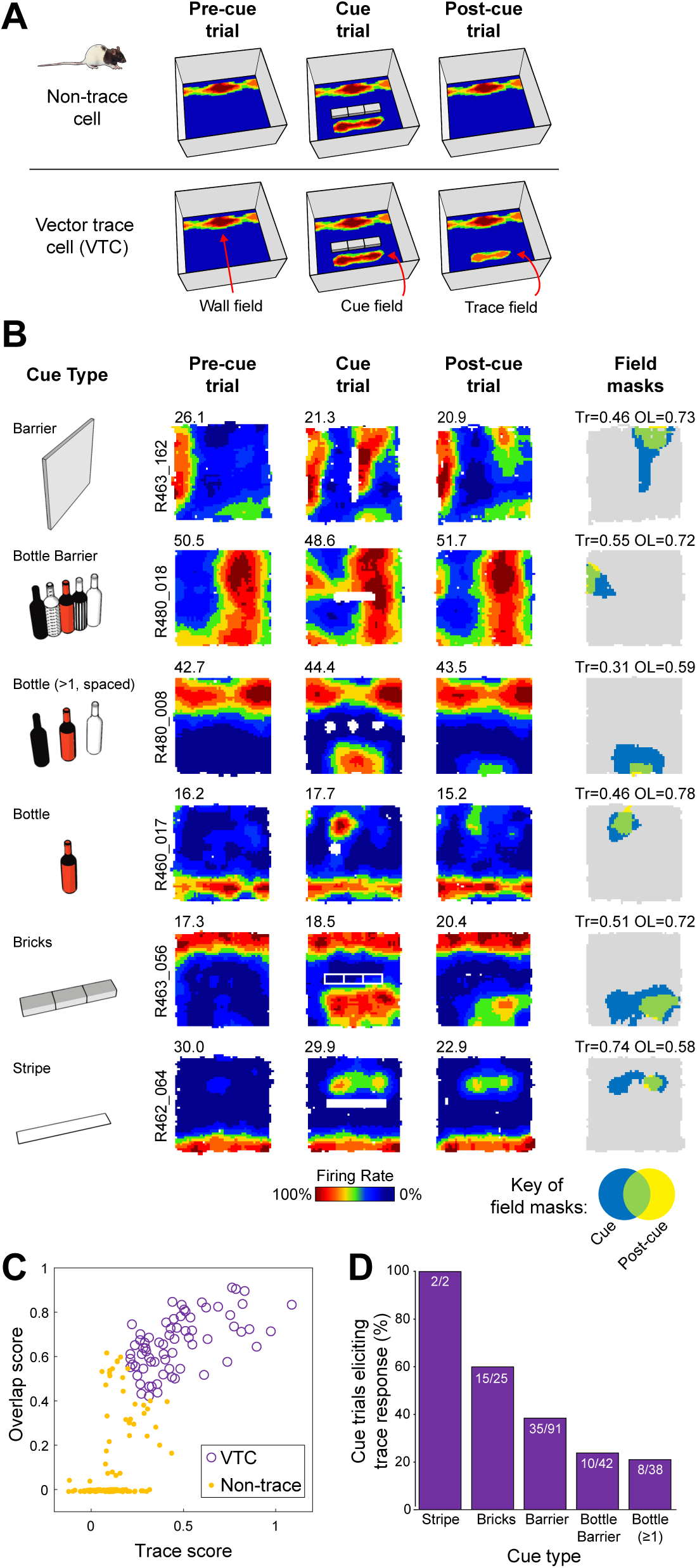
Vector trace responses to a range of cues. (**A**) Schematic of experimental procedure, including illustrative results. Rats foraged for food while neurons were recorded in the dorsal subiculum. Heat maps show firing rate of neuron as a function of rat’s position (warmer colours = higher firing). Top row: schematic of a ‘non-trace’ vector cell responsive to insertion of a new cue (‘Cue’ trial) but lacking a memory ‘trace field’ in the ‘Post-cue’ trial. Bottom row: a vector trace cell (VTC) whose cue-responsive firing field persists during ‘Post-cue’ trial following cue removal. (**B**) 6 representative VTCs. Left column: type of cue used. Middle columns: firing rate maps, peak firing rate (Hz) top-left of each map, rat and cell identifier numbers left of maps. Each cell (row) forms new firing field when cue (white space/lines) is introduced (Cue). Following cue removal (Post-cue), cue-responsive firing field persists in the region of the cue field. Right column: masks showing cue-responsive field (blue), post-cue field (yellow) and overlap between both, indicative of a trace response (green). Trace (Tr) and Overlap (OL) scores shown above each plot. (**C**) Scatter plot of Trace and Overlap scores for all neurons. VTCs are defined by combined above-threshold trace and overlap scores. (**D**) Percentages of tested cue-responsive neurons categorized as VTCs for each cue type. Fractions overlaid on bars show the number of cue responsive neurons recorded for cue type (denominator) and number of these neurons categorized as VTCs (numerator).

### Vector Trace cells respond to a wide range of cues, and exhibit longer distance tunings than non-trace cells

For vector coding to enable efficient navigation, it should be flexible, operating over a range of cue types and distances. Subiculum neurons were responsive to a range of different cue types (Fig. 1B, left column), and all cue-types were also capable of eliciting a memory-based response (Fig. 1D). Cue-responsive subiculum neurons are therefore capable of encoding allocentric vectors to a wide range of external cues, including both discrete objects and extended boundaries ^11^. We defined the distance tuning of vectors as the shortest distance between the peak firing bin of the cue-field and the edge of the inserted cue. VTCs’ distance tunings showed a larger variance and were on average longer than those of non-trace cells (Fig. 2A,B, Levene’s test p=0.0004; means-different t_196_ corrected=5.81: p<0.0001).

**Fig. 2.**
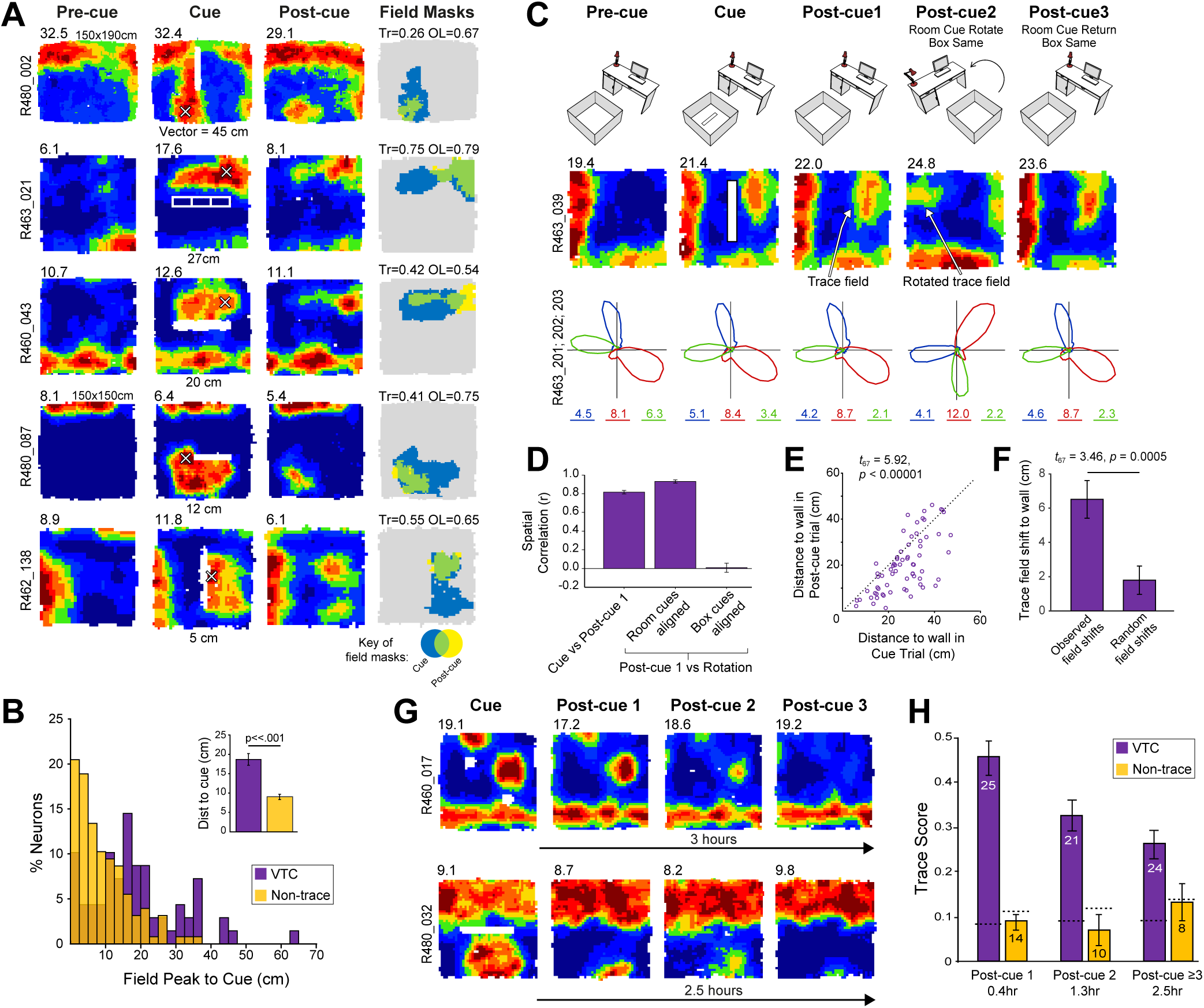
VTCs’ spatial field properties and hours-long persistence of trace fields following cue removal. (**A**) Rate maps of five VTCs (conventions as Fig.1) showing range of distance tunings (cue field-peak [white cross] to cue [white outline]). Vector length is shown below ‘Cue’ rate map. Non-standard box sizes are indicated above pre-cue rate map. (**B**) Larger variance and longer distance in VTCs’ vector tunings than those of non-trace cells. Inset: Mean (±SEM) vector tuning length VTCs and non-trace cells. (**C**) Room-cue rotation manipulation (room cues rotated 90° anticlockwise; box-and-floor unchanged). Top row: schematic of experiment. Middle row: rate map for representative VTC. Bottom row: polar firing rate plots showing directional tuning of three co-recorded head direction (HD) cells (marked in different colors, numbers below polar plots show peak rate). HD cells rotate with VTC cell. (**D**) Mean (±SEM) inter-trial correlations for VTCs subjected to rotation trials, following post-hoc counter rotation of rate maps to align to either room, or box-and-floor, cues. (**E**) Distance from cue and trace fields to the closest wall of the box, in cue- and post-cue trials. Each point shows the distances for one VTC. Trace fields systematically shift towards the closest wall, in the post-cue trial. (**F**) Mean (±SEM) trace field shift towards wall for observed data and simulated random instability of trace field (see Methods). (**G**) Hours-long persistence of trace fields in two representative VTCs in absence of cue. (**H**) Mean (±SEM) trace scores for VTCs recorded over multiple post-cue trials. Trace field strength in VTCs decays over time but remains significantly above chance; in non-trace cells it never exceeds chance levels (dashed horizontal lines; see Methods).

### Vector trace fields reflect memory, not responses to lingering odour cues

Could VTC trace fields reflect perceptual responses to odour cues left behind by objects rather than memory? Two lines of evidence speak against this possibility. Firstly, a sub-set of VTCs (19%, 13/70) in post-cue trials were subjected to the rotation of either the animal’s box-and- floor (‘intra-box’ cues) or room cues outside of the box (‘extra-box’ cues). In all cases, VTC field location was concomitant with extra-box, not intra-box cues (Fig. 2C,D, Fig. S2). When co-recorded, head-direction cells rotated in synchrony with VTCs (Fig. 2C, bottom row), suggesting that VTCs are integrated into global hippocampal mapping, rather than responding to discrete local odour cues. Secondly, in the post-cue trial, trace fields demonstrated a systematic shift away from the cue-trial position, and towards the nearest wall (Fig. 2E,F), suggesting an interaction between remembered and physically-present cues rather than a fixed response to a local cue. Taken together, these observations strongly argue that trace fields reflect mnemonic processing rather than responses to persistent local cues.

### Vector trace memory lasts for hours

To test how long VTC memory persists, a subset of VTCs were exposed to several post-cue trials in succession (Fig. 2G,H). Over 0.4-2.5 hours (2-4 exposures), mean Trace scores of VTCs were consistently higher, whilst those of non-trace cells were consistently lower, than chance (see Methods; Fig. 2H), demonstrating that VTCs encode memories of cue location over behaviourally-relevant timescales. The temporal decay in trace strength may additionally signal how long ago a cue was encountered.

### Proximo-distal axis: vector trace cells are common in distal subiculum but rare in proximal subiculum

The proximo-distal anatomical axis is considered a major organizational feature of the subiculum, based on patterns of connectivity ^17, 18, 25–29^, and gene expression ^18, 19^, with proximal subiculum thought to support *‘What?’* memory and distal subiculum *‘Where?’* (allocentric) memory ^18, 20, 30^. However, *in vivo* electrophysiological evidence for proximo-distal functional specialisation is largely absent ^11, 31^ consisting, at best, only of a gradient in mean firing rate and modest changes in spatiality of firing, e.g. ^32, 33^. However, here, strikingly we find that VTCs are overwhelmingly found in distal subiculum (Fig. 3; distal VTCs constituting 67/162 (41%), and proximal VTCs 2/34 (6%) of cue-responsive cells; n=196, χ^2^(1)=15.50, p=0.00008), strongly suggesting that distal subiculum has a specialised role in spatial memory.

**Fig. 3.**
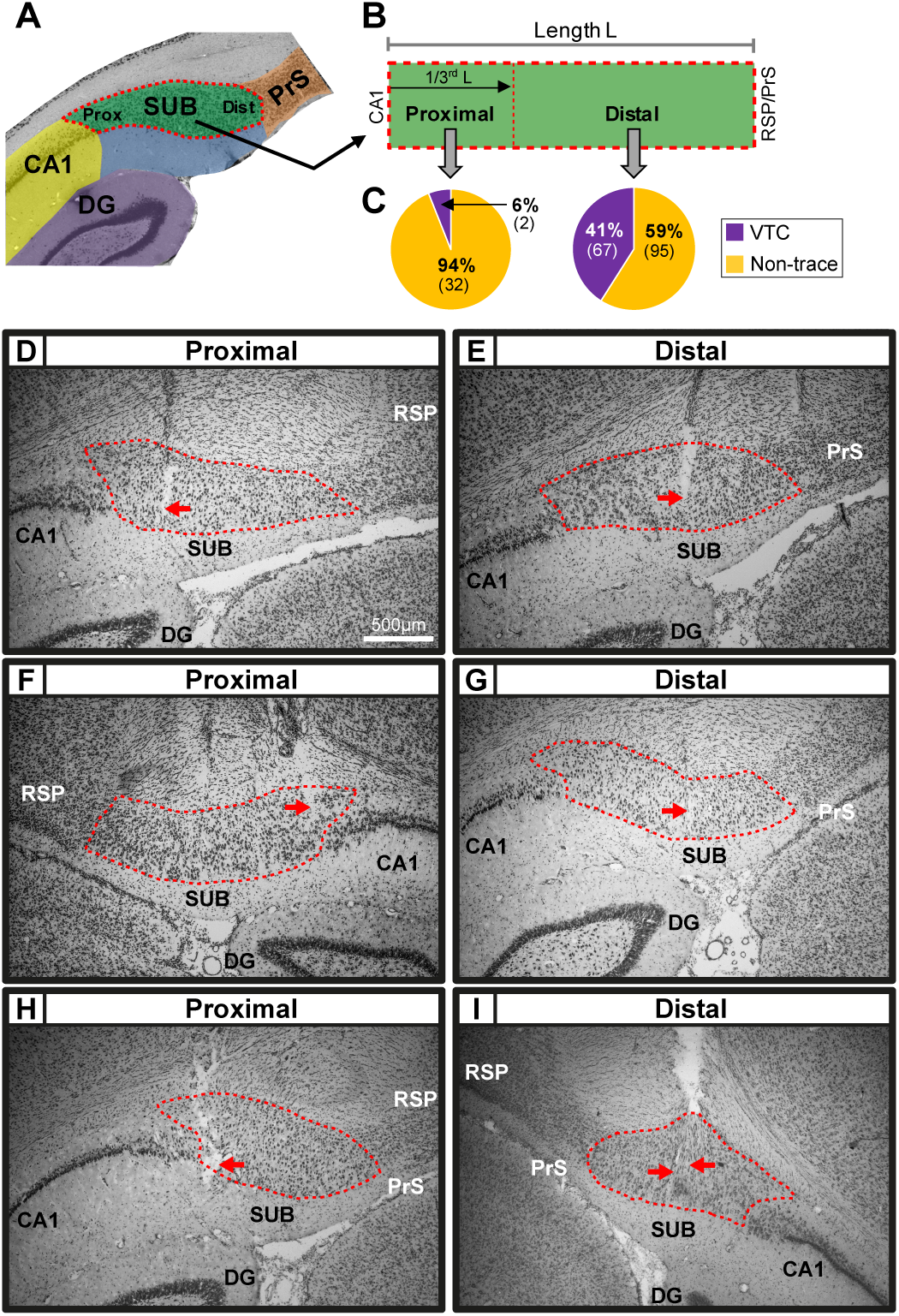
Vector Trace cells were common in distal subiculum, but rare in proximal subiculum. (**A** and **B**) Recording sites were assigned to proximal subiculum (the third nearest CA1), or distal subiculum (the two thirds furthest from CA1), divisions corresponding to previously reported connectivity and gene-expression patterns (see Methods). (**C**) High proportion of Vector Trace cells in distal, but not proximal, subiculum. (**D** to **I**) Photomicrographs of coronal sections depicting representative recording sites in proximal (D,F,H) and distal (E,G,I) regions. Red arrows point to estimated final location of the recording tetrodes. Dashed red lines indicate borders of subiculum’s pyramidal layer. Scale bar in (D) also applies to (E to I). Rat: 480 (D,E); 495 (F,G); 496 (H); 463 (I). Abbreviations: DG: Dentate gyrus; PrS: dorsal Presubiculum; RSP: Retrosplenial cortex; SUB: Subiculum.

### Proximo-distal axis: cue-responsive cells in distal division of subiculum fired at an earlier phase of theta than proximal cue-responsive cells

Spike timing with respect to the ongoing theta oscillation may modulate hippocampal neurons’ function in memory encoding and retrieval ^5, 34–36^. We therefore tested theta-modulation of cue-responsive neurons. We first tested whether the degree of theta modulation and preferred phase of cue-responsive neurons differed across the proximal and distal divisions of the Subiculum in the baseline situation. While theta modulation was similar across anatomical sub-divisions (Rayleigh r, pre-cue trial, distal: 0.16 ± 0.01; proximal: 0.14 ± 0.01; T_194_=1.22, p=0.22), distal cue-responsive cells fired at a considerably earlier phase of theta than proximal cue-responsive cells (Fig. S3; 87°, Watson-Williams *F*_1,190_=96.36, *p*<1×10^-12^).

### Theta phase indexed mismatch detection: preferred theta phase shifted in response to a novel cue but remained consistent in response to the familiar cue

To investigate theta-modulation of VTCs, we focused on distal subiculum, where the large majority of these cells were found (67 out of 69). Encoding-vs-retrieval scheduling models ^5, 34, 35^ predict that these memory processing states are separated neurally by the theta phase of spiking, with phase linked to direction (potentiation, depression) and strength of synaptic plasticity. Overall theta modulation of firing was similar for VTCs and non-trace cells (Rayleigh r, pre-cue trial, Trace: 0.15 ± 0.01; Non-trace: 0.16 ± 0.01; T_160_=1.00, *p*=0.32). Consistent with theta scheduling models, cue insertion elicited a markedly different (earlier) preferred phase of theta in the newly-generated cue-field in all cue responsive cells (Fig.4A, top row; distributions, statistics: Fig S4). Phase in the cue field (encoding) was earlier than in the same area in the pre-cue trial (baseline) and post-cue trial (retrieval) (VTCs: −35.1°, −36.2°; non-trace cells: −24.7°, −25.8°: all *p*<0.00012). Importantly, preferred theta phase in wall fields remained constant throughout the trial sequence (Fig 4A bottom row, Fig S5), ruling out that cue-field earlier phase is driven by altered global state. Thus, earlier phase indexed ‘mismatch’ detection ^1, 5^ which occurred in different box locations in a neuron-specific manner.

**Fig. 4.**
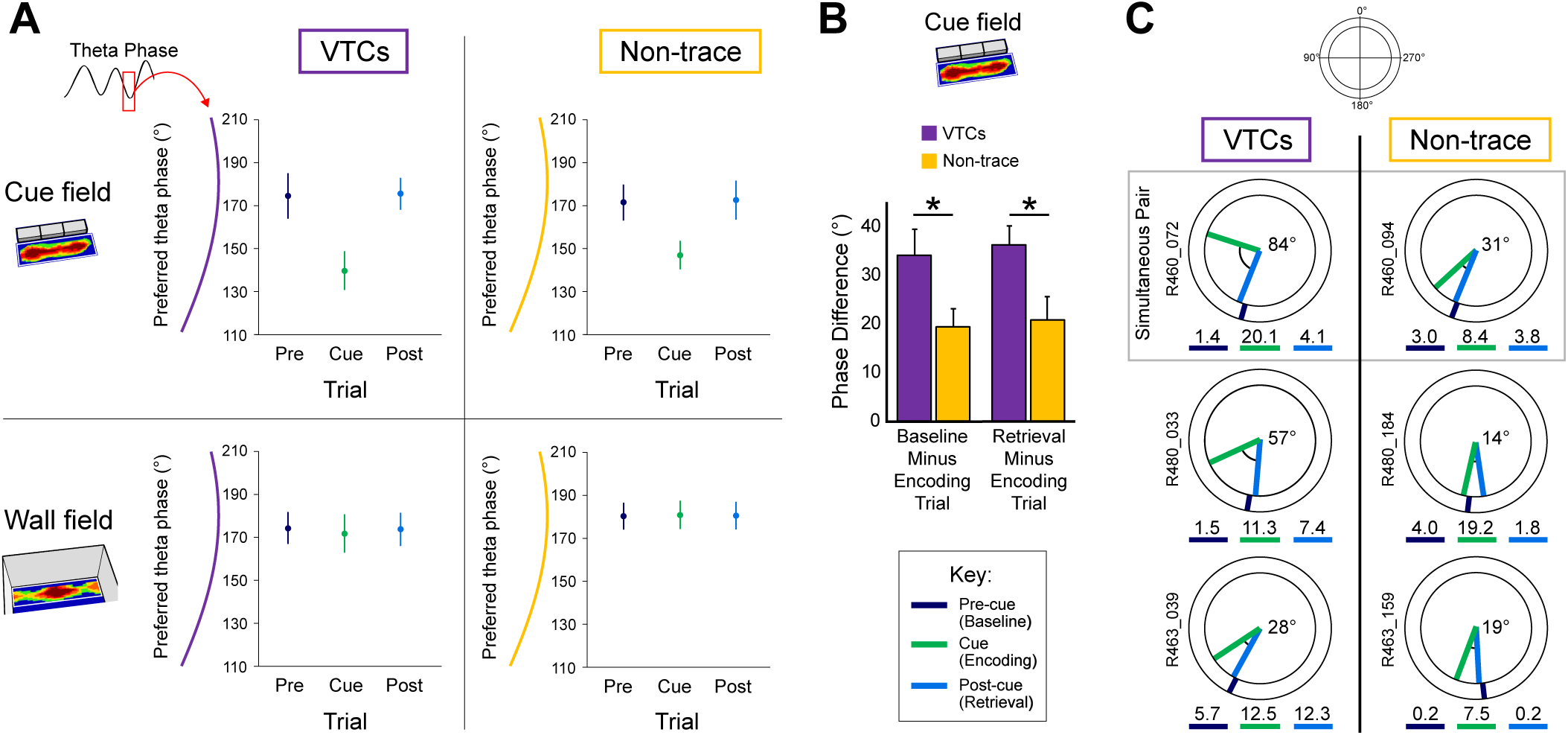
Earlier-going theta phase shift in encoding is greater in VTCs than non-trace cells. (**A**) Mean (±95% CI) preferred theta phase for neuronal firing, for the cue field (top row) and wall field (bottom row) areas. Preferred phase is markedly earlier in cue field in encoding trial, for both VTCs (left) and non-trace cells (right) (all Cue-vs-Pre-cue & Cue-vs-Post-cue *p*’s<1.2×10^-5^, see Fig.S4), but is stable for wall fields in all trials (all trial comparisons *p*’s >0.65, see Fig.S5); (**B**) Mean (±SEM) within-cell change in preferred phase in cue field area between baseline and encoding (left bars), and encoding and retrieval (right bars). Earlier-going phase shift, specifically in the encoding trial, is greater in VTCs, *: p=0.02. (**C**) Examples of preferred phase in in cue field area, in single neurons (top VTC and non-trace co-recorded pair) from three rats, with larger earlier-going shift in VTCs (shown in black at plot centers). Radial colored ticks show mean preferred phase in each trial, numbers below phase diagrams show mean firing rate (Hz) in each trial, see box for colour key.

### Cue-field theta phase change was greater in Vector Trace cells than non-trace cells

Models of hippocampal memory operations have long posited that hippocampus detects mismatches, and mismatch drives encoding, e.g. ^1, 5^. Could theta phase in the cue-field determine whether cue-responsive cells form a memory trace or not? There was no significant difference between the *absolute* phase of VTCs and non-trace cells in the cue-field (Fig S4). However, we also tested whether the amount of within-cell late-to-early phase change following the insertion of a cue was associated with forming a memory trace. We found that VTCs showed larger phase changes in the cue-field than non-trace cells, both when comparing cue-trial (encoding) to post-cue trial (retrieval) (Fig.4B, t_155_=2.35, p=0.02, Cohen’s *d*: 0.39), and cue-trial (encoding) to pre-cue trial (baseline) (Fig.4B, t_153_=2.31, p=0.02, Cohen’s *d*: 0.37). Fig.4C shows examples of individual cells. These results were not driven by large (and thus potentially circularly ambiguous) phase differences, since excluding the small number of cells that shifted preferred phase by more than 120° in either direction made the average effect size larger (*post-cue to cue*: n=152, Cohen’s *d*: 0.46, p=0.006; *pre-cue to cue*: n=151, Cohen’s *d*: 0.36, p=0.029). We emphasise that the changes in theta phase predicted by encoding-vs-retrieval models ^5, 34^ observed here do not occur in a particular region of space that is consistent for all cells, as in say the choice-arm (vs a side-arm) of an alternation maze ^35^. Rather, cue and trace fields were distributed over the entire environment, and indeed these cue-related fields for one cell will thus sometimes occur in the same region as the wall field of another cell (Fig. S6). Furthermore, these cell-type specific theta phase changes were not driven by changes in firing rate (Fig.S7).

## Discussion

In summary, our results identify a new category of neuron, the vector trace cell, defined by two properties. First, a VTC responds when the rat is at a specific distance and allocentric direction from a small or extended cue, including environmental boundaries, by immediately generating a vector field. Second, the vector field persists after the cue that elicited it is subsequently removed, creating a vector trace field.

Our findings build on an emerging picture of the prevalence of vector coding in the hippocampal formation, but add the crucial dimension of vector memory. Thus, neurons in the medial ^12^ and lateral ^37^ entorhinal cortex can encode allocentric vectors to discrete objects (though not environmental boundaries) but do not show memory for previously-present cues. Likewise, recent reports of egocentric vector coding of environmental cues ^38, 39^ also do not describe memory traces. Lateral entorhinal neurons ^40^, and a few CA1 place cells ^13^ can encode a memory trace for previously-present objects, but these non-vectorial object fields, which develop only after cue removal, are confined to the exact object locations so cannot retrieve locations in the space between objects and boundaries as is the case for VTCs. One study found 12% of CA1 bat cells showed *egocentric* vector tuning to a goal, some with memory responses ^41^. Interestingly, however, these goal-directed CA1 cells appear to encode the vector to only a single goal at a time. The present study is the first to report a cell class that combines encoding the location of multiple cues using allocentric vectors, with memory of those cue locations when the cues are removed. VTCs suggest a vector-based model of computing spatial relationships between an agent and multiple cues (or between different cues), freed from the constraints of direct perception of those cues, thus enabling spatial planning and imaginative cognition ^4, 7–9^.

The relative abundance and scarcity of VTCs in the distal and proximal subiculum, respectively, provide the strongest *in vivo* electrophysiological evidence for hypothesised spatial memory specialisation along the CA1/subiculum proximal-distal pathway ^30^, and provide a cellular substrate for the demonstration that selective inactivation of distal subiculum disrupts spatial memory ^18^.

Finally, we note that, consistent with encoding-vs-retrieval scheduling and dual-input control models of theta ^5, 34, 35^, all cue-responsive neurons encoded the presence of an inserted cue using an earlier phase of theta. The shift to earlier phase was specific to the cue-field, therefore not driven by global state changes nor, given the range of vectors especially in VTCs, restricted to a region in space. Rather, theta phase appeared to define a specific information channel for the presence of a newly-inserted cue within each neuron. Furthermore, the degree of relative late-to-early shift, within each neuron, was linked to whether a trace field would be formed. Thus our findings extend theta-scheduling models ^5, 34, 35^ by demonstrating theta-phase shift as a neural substrate for memory encoding. It will be important to determine the VTC-specific factors, such as particular anatomical inputs and/or enhanced plasticity, underlying VTCs’ theta-linked propensity to generate trace fields.

We suggest that given VTCs’ flexible responsiveness to diverse cues both within and at bounding perimeters, subicular VTCs constitute a powerful universal code for long-term vector memory in the hippocampal formation, of utility in spatial and likely non-spatial cognition.

## Acknowledgments

We acknowledge funding from the BBSRC (BB/M008975/1 to CL & co-I TJW); the Royal Society (UF100746 to TJW); the MRC (MR/N026012/1 to TJW) and start-up funds from CIMEC, Trento, Italy (SAL).

## Author contributions

CL, TJW and SAL contributed to funding acquisition. SP and CL conceptualised and designed the study. SP performed the experiments. SP, TJW and CL analysed the data and wrote the manuscript. SAL contributed to conceptualisation, experimental design and training. JD contributed to training, surgery and histology. All the authors discussed the results and contributed to the manuscript. We thank Neil Burgess and Daniel Bush for comments on an earlier version of this manuscript.

## Data and materials availability

Data and code are available upon reasonable request from the authors.

## Supplementary Materials

### Materials and Methods

#### Subjects

Six male Lister Hooded rats, weighing 392-522g at the time of surgery, were used as subjects. All rats were maintained on a 12:12 hour light:dark schedule (with lights off at 10:00am). Food deprivation was maintained during recording periods such that subjects weighed 85-90% of free feeding weight.

#### Surgery and Tetrode implants

Under deep anaesthesia (Isoflurane: initially at 3%, progressively getting lower to ∼1% thereafter) and using intra- and post-operative analgesia (Buprenorphine, 0.04mg/kg), rats were chronically implanted with two microdrives (one above the dorsal subiculum of each hemisphere). These microdrives allowed 4 or 8 tetrodes to be vertically lowered through the brain after surgery. The 8-tetrode loaded microdrives (implanted in four rats) used custom 3D-printed barrels to create a 4 × 2 tetrode array (150μm spacing between holes of barrel). With the 4-tetrode loaded microdrives (implanted in two rats), all the tetrodes were loaded using a single cannula. Tetrodes were constructed from HM-L coated platinum-iridium wire (90%/10%, California Fine Wire, 25μm). The details of tetrode mapping for each drive were recorded using photographs and notes both before surgery and after perfusion.

#### Subiculum: Implant co-ordinates and histology

Our implants targeted the anterior portion of the dorsal subiculum (i.e. near the septal end of the subicular septotemporal long axis). The skull coordinates used for insertion of the centroid of the tetrode array were based upon ^42^ in the following range: AP: −5.8 to −6.4 mm; ML: ±2.9-3.3 mm. Details of recording sites were reconstructed using records of electrode movement, physiological markers and post-mortem histology. The rats were killed and perfused transcardially with saline followed by 4% paraformaldehyde. Each brain was sliced coronally into 40-μm thick sections, mounted and Nissl-stained (using Cresyl-Violet or Thionin) for visualisation of the electrode tracks/tips. Data from recording sites in CA1 or dorsal presubiculum were excluded, and recording sites in subiculum were classified as being located in either the *Proximal* subiculum or *Distal* subiculum (see next section). Representative recording locations are shown in Figure 3.

#### Parcellating the subiculum into Proximal and Distal zones

The proximodistal axis has long been thought to be important in the functional anatomy of the subiculum, with the distal and proximal subiculum linked to spatial and object memory respectively (e.g. ^17, 20, 25–28, 30, 33, 43^. In the Subiculum, the Proximal pole abuts CA1, and the Distal pole abuts the Retrosplenial cortex/Presubiculum. In the present study, the coronal sections were used to classify recording sites as belonging to either the Proximal zone, consisting of the third closest to CA1, or to the Distal zone, consisting of the two-thirds furthest from CA1. The rationale for this approach was twofold. First, the Distal two-thirds of the Subiculum define a region that projects heavily to cortical regions linked to spatial memory (Medial Entorhinal, Retrosplenial, dorsal Presubicular and Parasubicular cortices; e.g. ^28, 44^. Second, the Proximal Subiculum (termed ‘Prosubiculum’ in ^19, 29^ as defined by gene and protein expression patterns, occupies approximately one-third of the Proximodistal extent of the anterodorsal Subiculum (see coronal sections #81-89 in Supplementary Figure 1 of ^19^; also see ^45^.

#### Electrophysiological recording

Rats were allowed a week to recover post-operatively before screening sessions began. During screening and inter-trial intervals, the rat rested on a square, holding platform (40cm sides, 5cm high ridges) containing sawdust. Tetrodes were gradually lowered towards the Subiculum pyramidal layer over days/weeks. Tetrodes were left to stabilize for least twenty-four hours after tetrode movement before recording commenced. Electrophysiological data from screening and recording sessions was obtained using Axona DACQ systems (DacqUSB). Electrode wires were AC-coupled to unity-gain buffer amplifiers (headstage). Lightweight wires (∼4 meters) connected the headstage to a pre-amplifier (gain 1000). The outputs of the pre-amplifier passed through a switching matrix, and then to the filters and amplifiers of the recording system (Axona, UK). Signals were amplified (6.5-14K) and band-pass filtered (500 Hz-7kHz). Each channel was continuously monitored at a sampling rate of 50 kHz and action potentials were stored as 50 points per channel (1ms, with 200 μs pre-threshold and 800μs post-threshold) whenever the signal from any of the 4 channels of a tetrode exceeded a given threshold. Local-field potential (LFP) signals were amplified 3.5-5K, band-pass filtered at 0.34-125 Hz and sampled at 250 Hz. Two arrays of infrared light-emitting diodes (LEDs), one array larger than the other for tracking discriminability, were attached to the rat’s head to track head position and orientation using a video camera and tracking hardware/software (DacqUSB, Axona, UK). The arrays of LEDs were positioned such that the halfway position between the two arrays was centred above the rat’s skull. Offline analysis defined this halfway position as the position of the rat (TINT, Axona, UK). Positions were sampled at 50Hz.

#### Testing laboratory and Recording Environments

External cues such as a lamp, PC monitor and cue cards on the walls provided directional constancy throughout the test trial series. For every trial, the rat was carried directly from the holding platform with its head facing towards the recording arena. During trials, the rat searched for grains of sweetened rice randomly thrown into the box about every 30 seconds. At the end of each trial, the rat was removed from the recording environment, and placed back on the holding platform until the next trial. Inter-trial intervals varied from 10 minutes to an hour but were typically around 25 minutes. The standard recording environment was a square box (100×100×50cm high) painted in ‘light rain’ grey. Occasionally, for more distally tuned cells, a larger, same coloured environment was used (either 150×150×50cm high or 150×190×50cm high). Four types of cue, introduced into the box during the ‘Cue trial’, were used (refer to Fig 1 for diagrams): a black painted barrier (50×2.5×50cm high); three wooden black bricks juxtaposed along their long axis (20×9.5×4.5cm high), thus creating an 60×28.5×4.5cm high cue; a high-contrast white stencil-painted stripe (60×10×0cm high) acting as a purely visual cue; and wine bottles (base diameter 7cm, 30cm high) painted with different large, high-contrast patterns and/or affixed with somatosensorily-different patches. In some trials, only one bottle was inserted into the environment. In other trials, two or more bottles were inserted in different configurations: tightly-juxtaposed in an array to create a continuous barrier; placed apart to create a linear spaced array; or placed apart in different regions of the box.

#### Standard test trial sequence and variants

The standard test trial sequence consisted of three consecutive trials: 1) a ‘pre-cue trial’ in which the recording box contained no cue; 2) a ‘cue trial’ in which the box contained one of the above-mentioned four types of cue; 3) a ‘post-cue trial’ in which the box again contained no cue. In a minority of sessions, one variant of the standard test sequence was that more than one cue trial was run successively before the post-cue trial. In this case, only the first cue trial was used to define the cue field and cue responsiveness of the neuron. In other cases, the experimental session was extended to include a repeat of the standard three-trial sequence (pre-cue trial, cue trial, post-cue trial), most often with physically different cues. In these cases, only the single cue trial with the strongest cue-elicited field (i.e. highest integrated firing rate, see description below), and its accompanying pre-cue and post-cue trials were selected for analysis.

#### Extended post-cue trial sequences including rotation trials

In some sessions, two or more post-cue trials were run successively. This enabled evaluation of how long trace fields persisted in the recording box in the continued absence of the cues that elicited them. In some instances, we also included a ‘rotation’ trial in this post-cue trial sequence to test for the potential influence of uncontrolled local odour cues upon trace fields. Rotation trials took two forms. 1) The local, intra-box cues (box-and-floor ensemble) was rotated 90° anticlockwise with respect to room. 2) External cues such as lamps and tables were rotated, while the intra-box cues maintained their orientation with respect to the testing room. As trace fields failed to rotate in all cases of type (1) rotation (see Figure 2), these trials were also included in the analysis evaluating how long trace fields persisted in the recording box, in the absence of cue objects.

#### Cell isolation

Cluster cutting to isolate single units was performed using a combination of Klustakwik v3 and manual isolation using TINT (Axona, UK). All the trials from a given session were loaded into Tint as a merged dataset, which was clustered using KlustaKwik’s principal component analysis. Subsequent manual adjustments were made where necessary. Once merged-trial cutting was complete, cell clusters were automatically split into each individual trial of that session (Axona MultiCutSplitter).

#### Firing Rate Maps

Firing rate maps for all recorded neurons were produced by first dividing the recording arena into a grid of 2×2cm square spatial bins, and finding the summed occupancy time and number of spikes fired in each bin. Summed occupancy and spiking maps were then smoothed with a 10×10cm boxcar kernel, and rate maps were constructed by dividing summed spiking by summed occupancy. Data from periods of immobility (movement speed <5cm/s) were excluded from rate maps.

#### Definition of cue fields and cue-responsive neurons

New firing fields generated by the insertion of a cue were detected as follows. First, firing rate maps were converted to z-scores, to allow comparison between different trials even following trial-to-trial fluctuations in firing rate. For each map, values for each bin had the overall mean firing rate subtracted, and were then divided by the overall variance of the firing rate across bins. Following this, the z-scored pre-cue map was subtracted from the z-scored cue map, thus producing a map describing where firing was increased specifically during the cue trial, relative to the pre-cue trial. Cue fields were then defined as contiguous regions of the resulting map with a value ≥ 1. If more than one cue field was present, only the largest was used for further analysis. Cue-responsive cells were defined as those where the sum of z-scored firing rate, within the cue field, was ≥70. Only recording sessions in which at least one VTC was recorded were included in the dataset, therefore the reported proportion of VTCs in the Subicular cue-responsive population is an upper-bound estimate.

#### Definition of Trace and Overlap scores

To define trace and overlap scores, cue and post-cue firing rate maps were z-scored, and the pre-cue map was subtracted from both, so as to highlight changes in firing relative to the pre-cue trial. (Same procedure as described above, ‘Definition of cue fields and cue-responsive neurons’). The Trace score was defined as the mean value of z-scored firing within the cue field region (as defined above) in the post-cue trial, divided by the mean of z-scored firing within the cue field region in the cue trial. A Trace score of 1 therefore indicates a memory-based response of equal strength to that induced by the presence of the cue. To define the Overlap score, we first detected whether any new regions of firing existed in the post-cue trial, relative to the pre-cue trial: such ‘post-cue’ fields were defined as contiguous regions of the (z-scored, pre-cue subtracted) post-cue map with value ≥1. If several post-cue fields existed, only the largest was used for further analysis. If a post-cue field was present, the Overlap score was defined as follows:

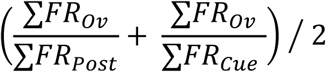

Where ∑*FR* in all cases refers to the summed firing rate in a region of the post-cue trial rate map: ∑*FR*_*Post*_ being the summed rate in the post-cue field, ∑*FR*_*Cue*_ the summed rate in the cue field, and ∑*FR*_*Ov*_ being the summed rate in the overlap between the cue and post-cue fields. Overlap score therefore assesses the average extent to which post-cue field firing overlaps the cue field firing, and vice versa. Where no post-cue field was present, the Overlap score was set to zero.

#### Spatially-randomised Trace and Overlap scores

To calculate whether the proportion of VTCs observed in our dataset exceeded that expected by chance, we constructed a population of Trace and Overlap scores derived from a spatially randomised dataset. Spatial randomization was performed by first calculating the cue field position from the cue map, as per normal analysis (see above). Then, the mask representing the region defined as the cue field was subjected to a random rotation and shift, before Trace and Overlap scores were calculated using post-cue map. Spatially-randomised scores therefore assessed the presence of memory-based firing, in a spatially random part of the post-cue trial. Spatial shifts of the cue field mask were subject to two constraints: 1) the shifted, rotated mask must lie entirely within the recording arena, 2) the shifted, rotated mask must not overlap with the pre-existing fields (‘Wall fields’) in the pre-cue trial (defined as contiguous areas with z-scored firing ≥ 1). This process was repeated 1000 times for each neuron. The proportion of spatial randomizations which resulted in a Trace and Overlap score that crossed our VTC classification thresholds (Trace ≥ 0.2, Overlap ≥ 0.4) was then calculated. The proportion of VTCs found in real data was tested against the proportion derived from spatially shuffled data, using a 1-sample Z-test.

#### Evaluating long-term trace field persistence in extended post-cue trial sequences

To test whether Trace scores during repeated post-cue trials exceeded chance levels, we used the same population of spatially-randomised Trace scores, generated as described above. To calculate if the mean of a given sample of Trace scores, of sample size N, was greater than expected by chance, we used the following procedure. N spatially-randomised Trace scores were randomly sampled (with replacement) from the random population, the mean of this sample was calculated, and the process repeated 100,000 times. The 95^th^ percentile of the resulting population of spatially-randomised sample means was defined as the 95% confidence level for the real Trace score mean to be greater than chance.

#### Movement of trace fields towards nearest wall

To measure the movement of memory-based responses towards the recording box walled perimeter, the post-cue field, rather than the cue-field region, was used to define a ‘trace field’ in the post-cue trial, so as to allow an unbiased estimate of whether memory-based firing is subject to spatial displacements between trials. We estimated the position of both the cue and post-cue fields, using the firing rate-weighted centroid of their respective field masks. The distance between the cue, and post-cue fields, and the closest wall, were then calculated. As inserted cues were, on average, situated towards the environment centre, even a random drift of field positions would have created an overall mean shift in field position towards the recording box walls. To control for this possibility, we constructed a spatially-randomised set of field shifts for each cell, and compared the overall mean shift towards the nearest wall for real and randomised data. Spatially random shifts were defined as shifts of the cue field, where the shift direction was random, and the shift distance was randomly drawn from a population matching the overall distribution of shift distances in the real data. Shifts were constrained such that the randomly shifted field needed to lie entirely within the recording box boundaries. 1000 random shifts were produced for each cell, and the mean random shift towards the nearest wall was calculated for each cell. The difference between real and mean random shifts was tested using a paired t-test.

#### Theta firing phase analysis

Instantaneous theta phase was defined by filtering LFPs using a 5-11Hz bandpass filter, and taking the angle of the Hilbert-transformed filtered signal. The theta phase of each spike was defined as the phase of the temporally corresponding LFP sample, from the hemisphere- and anatomical region- (proximal vs. distal subiculum) matched LFP with the highest signal-to-noise ratio for the theta oscillation. The signal-to-noise ratio for theta was defined as the mean power in the theta band (±0.5Hz around the highest power between 7-10Hz) divided by the mean power in the range 2-20Hz, excluding the theta band. Spectral power was estimated using the fast-fourier transform. The overall theta modulation of each neuron was estimated using the length of the resultant mean vector of phases. Only neurons with significant phase modulation (defined as Rayleigh test p<0.01) were used for phase analysis. The preferred firing phase of each neuron, in each firing field, was defined as the circular mean of the spike phases, for spikes occurring whilst the rat was within the given firing field (Wall field, Cue field regions). Distributions of neuronal preferred phases (e.g. Distal cells, VTCs) were characterized by: the mean phase, i.e. the circular mean of the neuronal preferred phases, 95% confidence limits, Rayleigh vector r, Von Mises κ (indexing concentration), and the circular standard error of mean (see Supplementary Figures). The difference between mean *absolute* preferred phases was tested using Watson-Williams F tests. The *shift* of preferred spike phase in the cue field induced by cue insertion was defined for each cell as the signed shortest circular distance between the mean phases in the cue trial, and either the pre or post-cue trial. The difference between VTC and non-trace cell phase shifts was compared using an independent samples t-test. As outlined in the main text, to control for the influence of large (and thus potentially circularly ambiguous) shifts, this analysis was repeated using only shifts of ±120°.

---

**Fig. S1.**
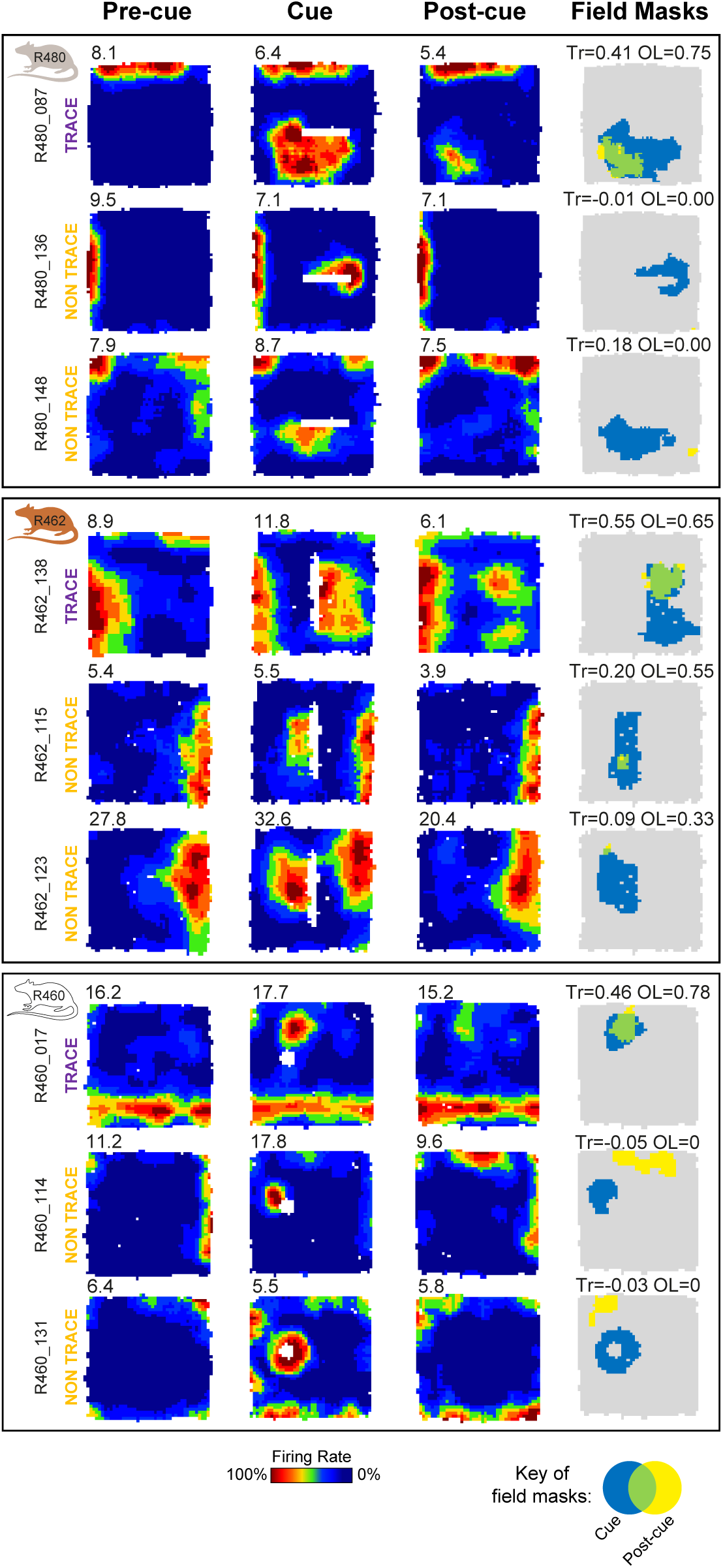
Co-recorded vector trace cells and non-trace cells. Firing rate maps of simultaneously-recorded vector trace cells and non-trace cells in three rats. Conventions as in Figure 1. Each row illustrates firing rate maps for one cell (peak-rate bin (Hz) top-left). **Left column**: Pre-cue trial. **Left-middle column**: Cue trial, where each cell forms a new firing field when the cue (white space) is introduced. **Right-middle column**: Post-cue trial. In vector trace cells (top row for each rat), following cue removal, the cue-responsive firing field persists. In contrast, in non-trace cells, the cue-responsive firing field has diminished to below-threshold levels in the Post-cue trial. **Right column**: masks showing cue-responsive field (blue), post-cue field (yellow) and overlap between both (green). Trace (Tr) and overlap (OL) scores shown above each plot. Trace score value of non-trace cell R462_15 is 0.198, i.e. below Trace cell threshold of 0.20.

**Fig. S2.**
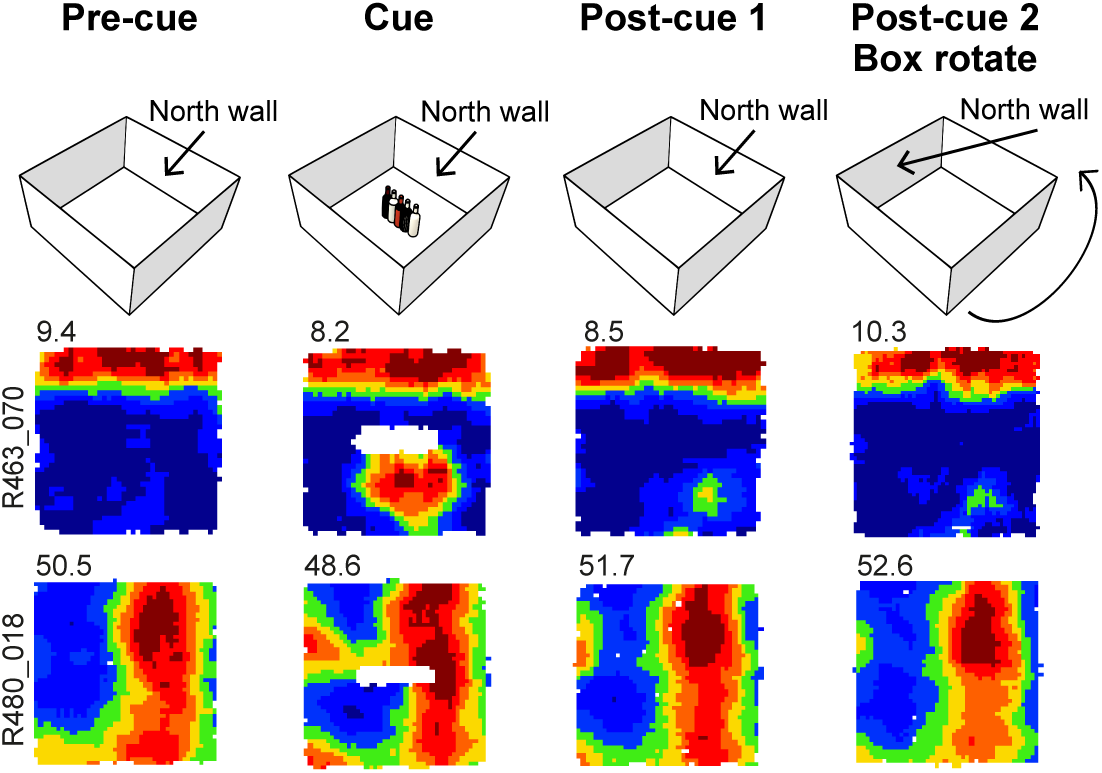
Rotational manipulations of intra-box cues show that trace fields do not reflect responses to local odour cues. Two example VTCs showing how rotating the walls and floor of the box has no effect upon location of VTC trace fields. **Top row** indicates sequence of four successive trials with two Post-cue trials. For Post-cue trial 2, box-and-floor configuration has been rotated by 90° anti-clockwise. **Middle row** (Rat 463) and **bottom row** (Rat 480): representative VTCs showing that the wall fields and trace fields are unaltered by the box-and-floor rotation.

**Fig. S3.**
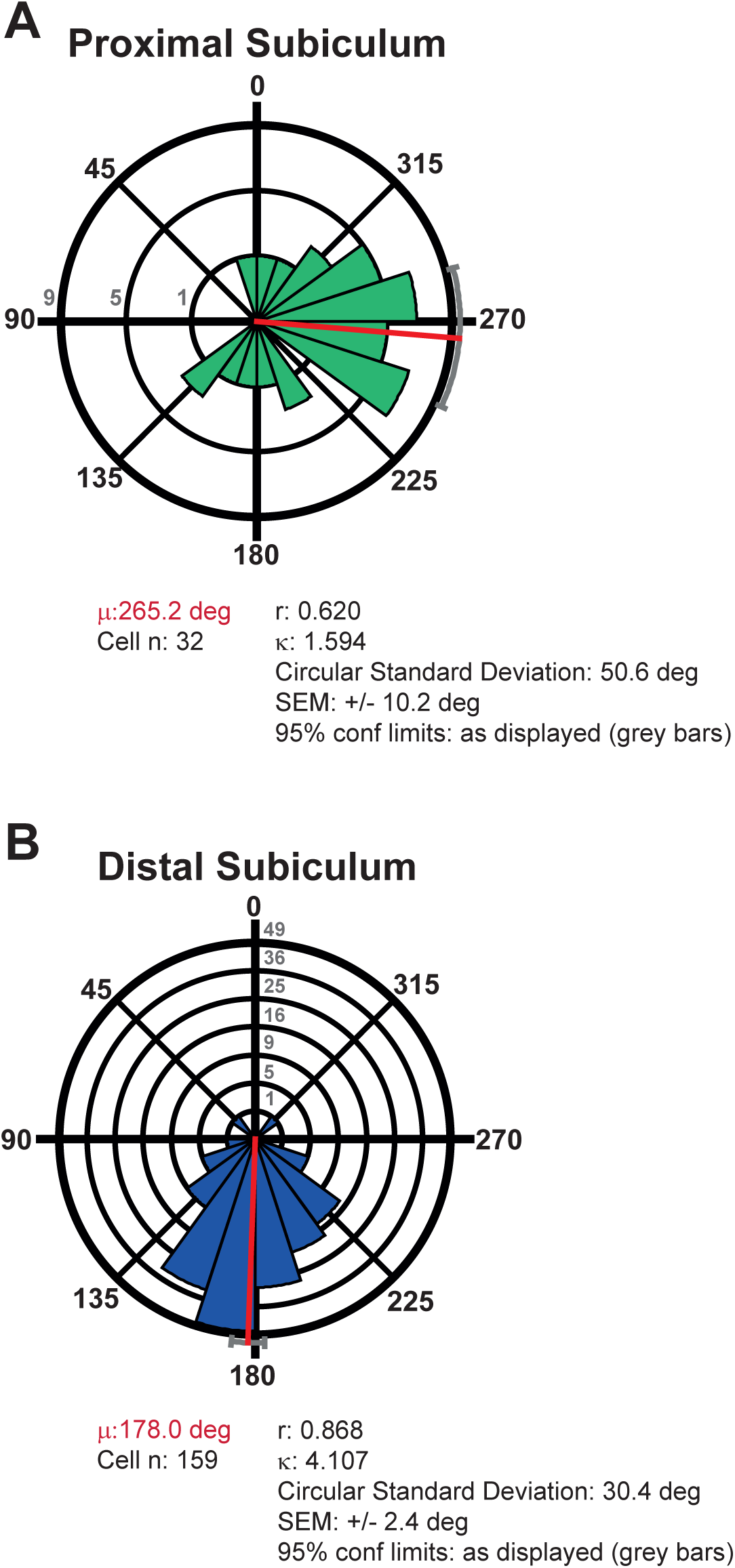
Preferred theta phase of firing in the distal subiculum occurs a quarter of a cycle earlier than in the proximal subiculum. (**A** and **B**) Polar histograms showing distribution of individual cell phases in the wall field area in the Baseline (Pre-cue) trial in the proximal subiculum **(A)** and in the distal subiculum **(B)**. Phase distribution is more concentrated in the distal subiculum, where mean preferred phase occurs a quarter-cycle earlier than in the proximal subiculum (87 degrees earlier: Watson-Williams *F* test for difference between means: *F*_1,190_=96.36, *p*<1×10^-12^). Phases are divided into twenty 18-degree bins (0/360=peak;180=trough). All cells included where Rayleigh test *p*<0.01. Scale near 90-degree line (A), and zero-degree line (B) indicates number of cells in a given phase bin. Key: µ is mean phase (red line, red font text), r is length of Rayleigh vector, κ is Von Mises’ κ (indexing phase concentration). Grey bars depict +/− 95% confidence limits.

**Fig. S4.**
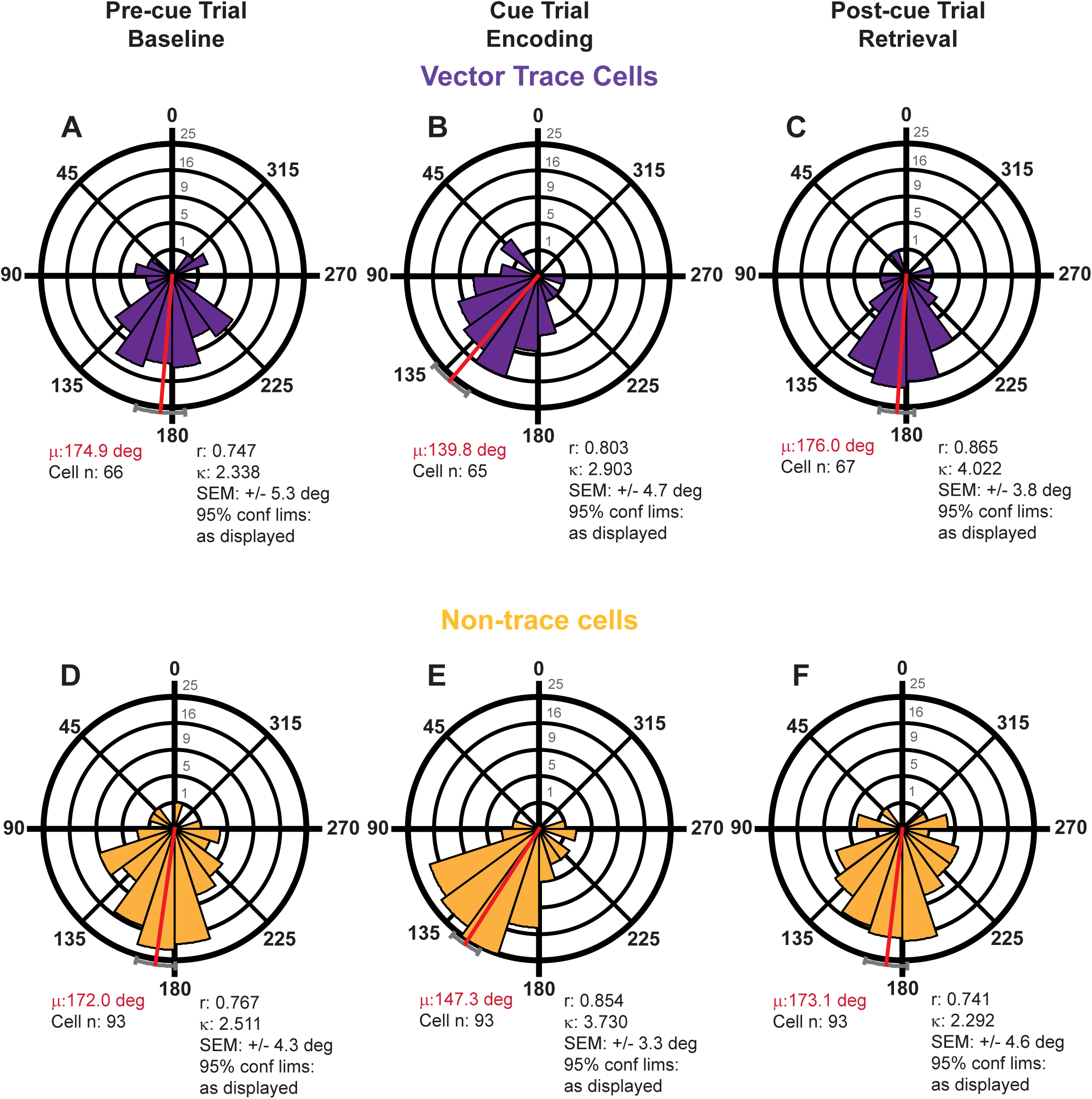
Preferred theta phase of firing shifts markedly earlier during cue-related encoding in both vector trace cells and non-trace cells. Polar histograms showing distribution of individual cell phases in the cue field area whose mean phases (here, red lines, red-font text) and 95% confidence limits (here, grey bars) are shown in Figure 4, for each of the two cell types: (**A** to **C**) trace cells, (**D** to **F**) non-trace cells, from Pre-cue trials (A,D), to Cue trials (B,E), to Post-cue trials (C,F). Preferred theta phase of firing is very stable across Pre-cue and Post-cue trials in both cell types. Notably, against this background of strong phase stability, preferred theta phase of firing is markedly earlier in both cell types during Encoding trials (Cue Trial: B,E) than during Baseline (Pre-cue Trial: A,D) and Retrieval (Post-cue Trial: C,F) trials (all Cue-vs-Pre-cue & Cue-vs-Post-cue *F* values >20.4, all *p* values <1.2×10^-5^). Within-cell earlier shift in Encoding was greater in vector trace cells than non-trace cells (main text; Fig 4.) There were no within-trial, across-cell-type, absolute-phase differences (Watson-Williams *F* tests: Pre-cue trial: *F*_1,157_=0.19, *p*=0.67; Cue-trial: *F*_1,156_=1.75, *p*=0.19; Post-cue trial: *F*_1,158_=0.22, *p*=0.64). Phases are divided into twenty 18-degree bins (0/360:peak,180:trough). All cells included where Rayleigh test p<0.01. Vertical scale near zero-degree line indicates number of cells in a given phase bin. Key: µ is mean phase (red line, red font text), r is length of Rayleigh vector, κ is Von Mises’ κ (indexing phase concentration). Grey bars depict +/− 95% confidence limits.

**Fig. S5.**
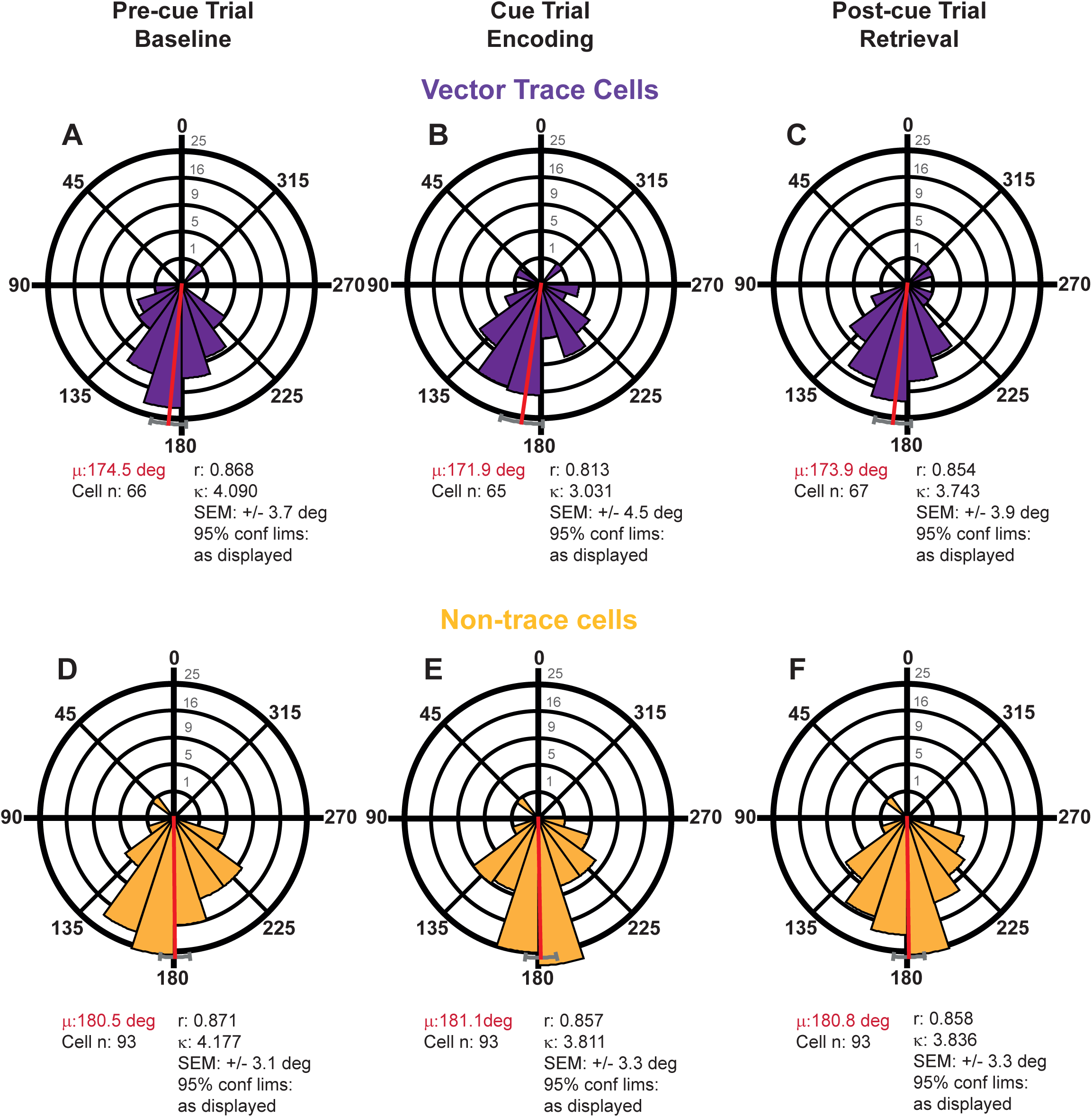
Preferred theta phase of firing is highly stable across trials in the wall field in both vector trace cells and non-trace cells. Polar histograms detailing distribution of individual cell phases in the wall field area whose mean phases (here, red lines, red-font text) and 95% confidence limits (here, grey bars) are shown in Figure 4, for each of the two cell types: (**A** to **C**) trace cells, (**D** to **F**) non-trace cells, from Pre-cue trials (A,D), to Cue trials (B,E), to Post-cue trials (C,F). Across-trial preferred theta phase of firing is very stable (Watson-Williams *F* tests comparing within-cell-type, across-trial, phase distributions: 2×3=6 tests; all 6 test *F* values <0.20, all *p* values >0.65). There were no within-trial, across-cell-type differences (Pre-cue; *F*_1,157_=1.52, *p*=0.22; Cue: *F*_1,156_=2.82, *p*=0.10; Post-cue: *F*_1,158_=1.84, *p*=0.18). Phases are divided into twenty 18-degree bins (0/360=peak; 180=trough). All cells included where Rayleigh test *p*<0.01. Vertical scale near zero-degree line indicates number of cells in a given phase bin. Key: µ is mean phase (red line, red font text), r is length of Rayleigh vector, κ is Von Mises’ κ (indexing phase concentration). Grey bars depict +/− 95% confidence limits.

**Fig. S6.**
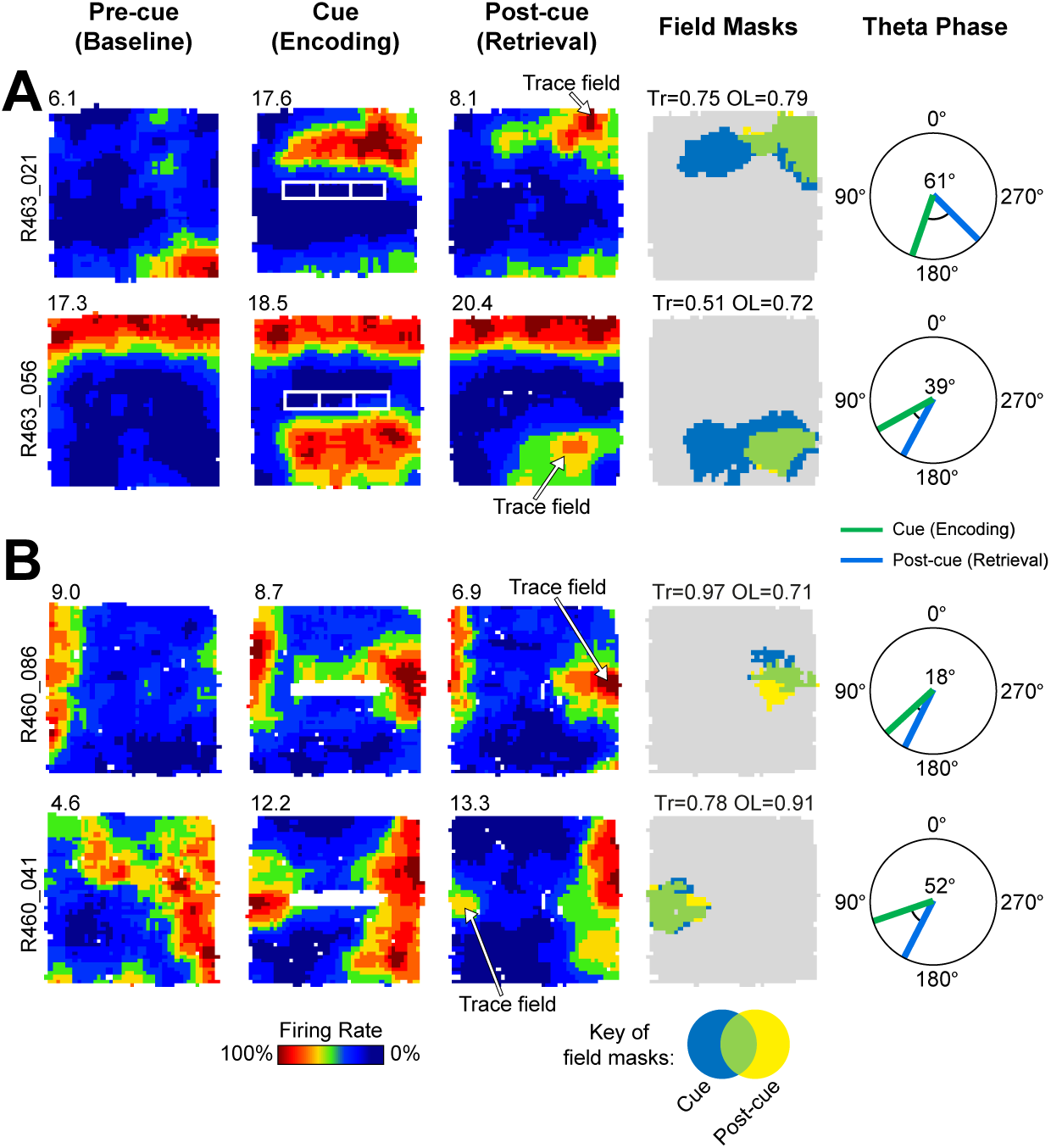
Examples of simultaneously-recorded vector trace cell pairs, demonstrating phase changes are specific to cue-driven firing, and not linked to one region in the recording arena. (**A** and **B**) Simultaneously-recorded pairs of vector trace cells in two rats. Columns left to right: rate maps for each cell across trials (pre-cue, cue, post-cue), field masks (cue and post-cue), and preferred (mean) theta phase of firing in Encoding and Retrieval trials (‘Theta phase in cue field region’). These examples show that the same cue (A: 4.5cm high brick array; B: 50cm high wall) elicits, in simultaneously-recorded cells, trace fields occupying different areas of the box, far from each other. VTC firing at earlier theta phase is specifically linked to the cue-driven firing field for each cell, and, across cells, is dissociated from the rat’s position.

**Fig. S7.**
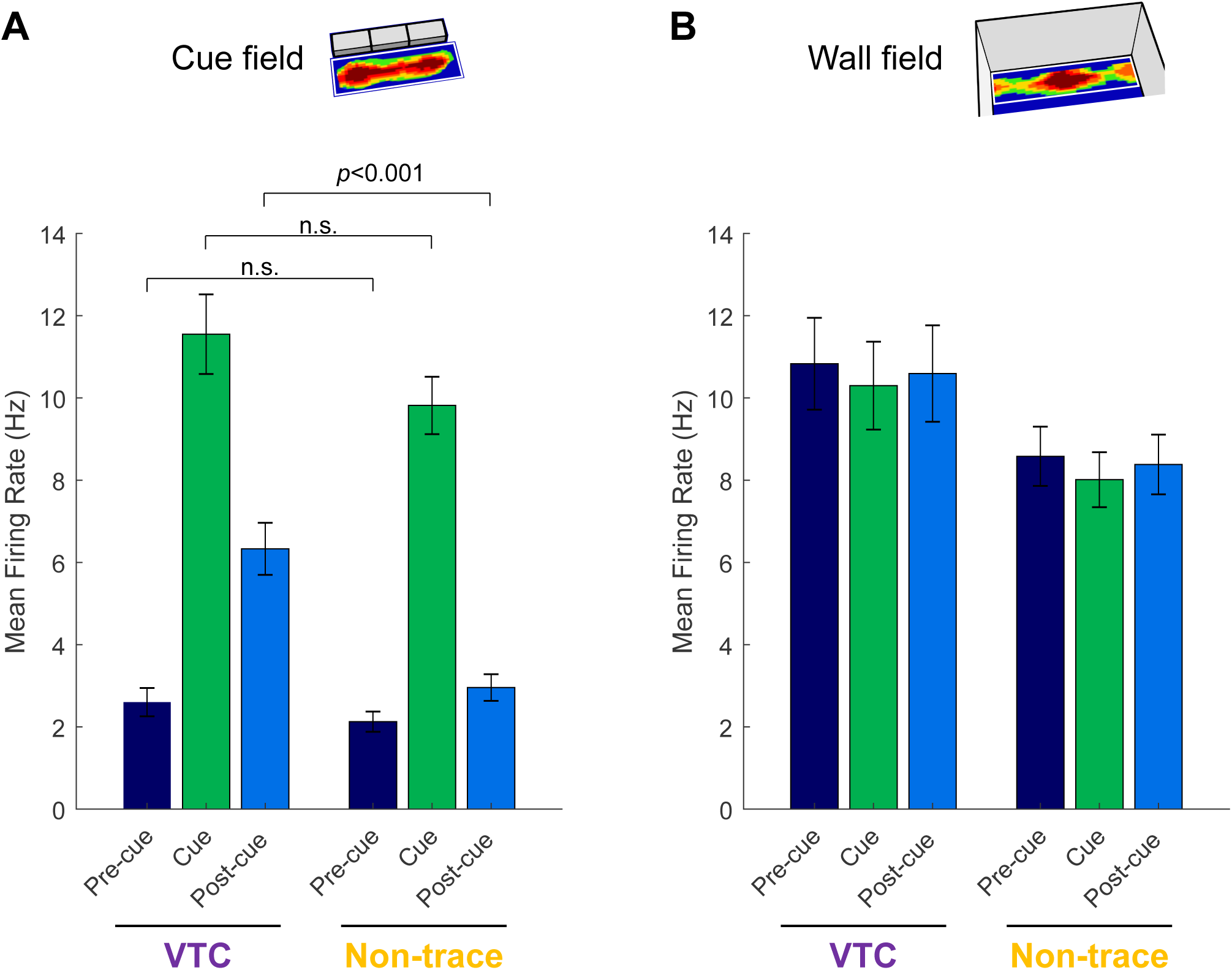
Firing rates in VTC and non-trace cells in distal subiculum. Bar graphs showing means (±SEM) of mean neuronal firing rates in the cue field (**A**) and wall field (**B**), in both VTCs and non-trace cells in distal subiculum. One possible account of changes in preferred phase (e.g. following cue insertion), is that these may reflect high-rate and low-rate regimes of firing respectively, under models of theta phase precession in which higher depolarisation drives earlier phase of firing ^46, 47 but see,48^. Here we show that, by contrast, phase and firing rate can be dissociated, in particular in the post-cue trial. Consistent with our cell-type classifications, VTC firing rates were significantly greater than non-trace cell firing rates in the cue field area, specifically in the post-cue trial (A: 2-way mixed ANOVA cue field firing rates: Trial*Cell type; F_2,304_=9.7 p<0.001; Post-hoc Simple Main Effects VTC vs Non-trace: Pre, p=0.34; Cue, p=0.11; Post, p<0.001). By contrast, there was no difference in the preferred phase of VTC and non-trace cells, in any trial, including the post-cue trial (see Fig 4, Fig S4, Watson-Williams *F* tests: Pre-cue trial: *F*_1,157_=0.19, *p*=0.67; Cue-trial: *F*_1,156_=1.75, *p*=0.19; Post-cue trial: *F*_1,158_=0.22, *p*=0.64). There is therefore no significant difference between VTCs and non-trace firing rate in the cue trial which could explain the greater earlier-going phase shift. Furthermore, there is a strong dissociation between phase and rate in the post-cue trial, whereby VTC rates are significantly higher than non-trace cell rates, but preferred phase is strongly similar for the two classes of neuron. In the wall field (B), VTCs showed a strong trend towards greater firing rates overall (2-way mixed ANOVA wall field firing rates: Trial; F_1,152_=3.85 p=0.052), but no cell-type specific changes in firing across trials (Trial*Cell type; F_2,304_=1.33 p=0.26).

